# Comparative genomics-based insights into diversification and bio-protection function of *Xanthomonas indica*, a non-pathogenic species of rice

**DOI:** 10.1101/2023.04.15.537001

**Authors:** Rekha Rana, Vishnu Narayanan Madhavan, Ramesh V. Sonti, Hitendra K. Patel, Prabhu B. Patil

## Abstract

*Xanthomonas* species have been extensively studied as major and model pathogens of plants. However, there is an increasing recognition of the complex and large world of non-pathogenic species of *Xanthomonas* in the recent decades. One pathogenic *Xanthomonas* species has been known in rice for the last hundred years, yet in recent years, three non-pathogenic *Xanthomonas* (NPX) species have been reported. Thus, there is a need to understand the species-level diversity of NPXs like their pathogenic counterparts. In the present study, we report isolation and investigation into the genomic diversity of *Xanthomonas indica*, a newly discovered NPX species from rice. The study allowed us to establish the relationship of *X. indica* strains within clade I of Xanthomonads. All the strains formed a distinct but diverse clade compared to clades corresponding to other NPX species from rice and other hosts. *X. indica* lacks major virulence factors found in their pathogenic counterparts. Identification of highly diverse strains and open-pan genome indicates ongoing selection to acquire new functions for adaptation. *X. indica* also harbours large-scale interstrain variations in the lipopolysaccharide O-antigen biosynthetic gene cluster, which hints at the selection at this locus. Interestingly, all the diverse strains of *X. indica* were able to protect rice from bacterial leaf blight pathogen of rice upon leaf clip inoculation. Comparative genomics revealed the association of a RiPP, a metalloprotease and a bacterial-killing type IV secretion system conserved across its related member species, including *X. sontii*, in the clade I with *in-vivo* anti-pathogenic properties. Overall the study has provided novel evolutionary insights into an important NPX member species. The findings and genomic resources will allow further systematic molecular studies to understand its interaction with the host plant.

## Introduction

The genus *Xanthomonas* comprises a large group of bacterial pathogens infecting a wide range of economically important plants (Ryan et al., 2011). There are thirty-five validly published species in the genus *Xanthomonas* (Parte, 2018)(www.bacterio.net). *Xanthomonas* genus is mainly known for the prominent phytopathogens associated with various cereals, fruits, vegetables, and ornamental plants. Recently, there have been increasing reports of non-pathogenic *Xanthomonas* (NPX) species being isolated from the same hosts on which their pathogenic counterparts cause diseases (Bansal et al., 2021; Bansal, Kumar, & Patil, 2020; Cesbron et al., 2015; Martins et al., 2020; Rana et al., 2022). So far, three NPX species have been isolated from rice plants alone, indicating an undiscovered world of *Xanthomonas* species with a non-pathogenic lifestyle (Bansal et al., 2021; Rana et al., 2022; Triplett et al., 2015). The NPX isolates are mostly ignored, for they lack any importance in the pathogenicity context. Nevertheless, their existence might serve some important functions to the host plants. Interestingly, recent studies have elucidated the bio-protection potential of these non-pathogenic strains against disease causal organisms of their hosts. *Xanthomonas sacchari* R1, a misclassified NPX strain from rice, is reported to show antagonism against *Burkholderia glumae*, the causal organism of panicle blight of rice (Fang et al., 2015). Similarly, another *Xanthomonas* strain isolated from perennial ryegrass showed strong bio-protection activity against fungal pathogens of grasses (Li et al., 2020). This relatively understudied members of the *Xanthomonas* community is likely to be significant for evolutionary studies and might also be involved in competitive relationships with the pathogenic residents of their hosts. The NPX could therefore be investigated for biocontrol potential against host plant diseases.

Rice is a staple food crop for half of the world’s population. *Xanthomonas oryzae* pv. *Oryzae* (Xoo) is the causal agent of bacterial leaf blight, a major disease in rice. Its variant pathovar *X. oryzae* pv. *oryzicola* (Xoc) is the causal agent of bacterial leaf streak disease in rice, and another pathovar from the species is *X. oryzae* pv. *leersiae* (Xol) (Lang et al., 2019). Interestingly, three NPX species have been reported from rice. First is *X. maliensis*, and the other two reported by our group are *X. sontii* and *X. indica* (Bansal et al., 2021; Rana et al., 2022; Triplett et al., 2015). This suggests the importance of the NPX community from rice. Although the pathogenic *Xanthomonas* species and pathovars are well studied at the population and functional level, there is a need to investigate strain-level variation in rice NPX species. In the same manner in which the rice-*Xoo* system has been an excellent model for plant-pathogen interaction, rice-NPX can be a promising model for understanding plant-microbe interaction studies. In this direction, the present study reports the isolation of genomically diverse strains of *X. indica* and insights into the evolution of this NPX species. Such a study allowed us to establish the phylogenetic relationship among NPX species of rice and other plants. The study also allowed understanding the sub-clades in clade I of Xanthomonads consisting of several pathogenic and NPX species/strains. Using comparative genomics, we also provide insights into the function and uniqueness of this NPX species as an important member of the rice microbiome. Further, we also provide evidence for *in-vivo* plant protection properties of *X. indica* and the evolution of this trait in other NPX species of rice and its relatives. The genomic analysis revealed the association of a metalloprotease and a bacterial-killing type IV secretion system (X-T4SS) in *X. indica* and its phylogenetically close relatives with respect to in-vivo protection of rice from *Xoo*. The resources and findings from this study will aid in further systematic molecular and functional studies of this important NPX species of rice.

## Material and methods

### Whole genome sequencing

Bacterial isolates PPL121, PPL129, PPL133, PPL134, PPL135, PPL139, PPL116, and PPL405, were isolated from rice seeds obtained from different geographic locations in India during the year 2021 as per previously described protocol (Cottyn et al., 2001; Midha et al., 2016). Genomic DNA was extracted using Quick-DNA™ Fungal/Bacterial Miniprep Kit (Zymo Research, USA). Genomic DNA quality and quantity assessment was carried out using Nanodrop 1000 (Thermo Fisher Scientific) and Qubit 4 fluorometer (Invitrogen). Whole genome sequencing was done using the Illumina Nova-seq platform facility (MedGenome, Hyderabad). Raw reads were subjected to quality assessment using FastQC v0.11.9 (https://qubeshub.org/resources/fastqc). Sequenced paired-end reads were assembled using SPAdes v3.13.0 and SPAdes v3.15.5 (Prjibelski, Antipov, Meleshko, Lapidus, & Korobeynikov, 2020). The genome assembly parameters were estimated using Quast v5.0.2 (Gurevich, Saveliev, Vyahhi, & Tesler, 2013). Assembled genomes were checked for completeness and contamination using CheckM v1.2.2 (Parks, Imelfort, Skennerton, Hugenholtz, & Tyson, 2015). Draft genomes were submitted to the NCBI GenBank database and annotated using the NCBI’s Prokaryotic Genome Annotation Pipeline (PGAP) (https://www.ncbi.nlm.nih.gov/genome/annotation_prok/).

### Phylogenetic and taxonomic analysis

Phylogeny was constructed using genomes of *X. sacchari* and *X. sontii* from the NCBI GenBank, along with other previously reported non-pathogenic strains. Type strains of some important *Xanthomonas* species were also considered for the phylogenetic analysis. 55 genomes were annotated using Prokka v1.14.6 to generate GFF files (Seemann, 2014). Roary v3.12.0 was used to create a concatenated core gene alignment using the GFF files as input at an identity cut-off of 90% (Page et al., 2015). The phylogeny was constructed using PhyML v3.3.20211231 with GTR model of phylogeny construction running 1000 bootstrap replicates (Guindon et al., 2010). The average nucleotide identity (ANI) values were calculated using oANI algorithm with USEARCH v11.0.667, and digital DNA-DNA hybridization (dDDH) values were calculated using formula 2 of web based Genome to Genome Distance Calculator v3.0 (I. Lee, Ouk Kim, Park, & Chun, 2016; Meier-Kolthoff, Carbasse, Peinado-Olarte, & Göker, 2022). CSI phylogeny version 1.4, available at the Centre for Genomic Epidemiology, an open web server, was used for SNP calling and phylogeny construction (Kaas, Leekitcharoenphon, Aarestrup, & Lund, 2014). *X. indica* PPL560^T^ was taken as a reference genome.

### Pangenome analysis

The pangenome of *X. indica* species was estimated using Roary v3.12.0 with BLASTp identity set at 90% using Prokka v1.14.6 generated GFF files as input (Page et al., 2015; Seemann, 2014). The pan-genome profile for the species was generated using PanGP v1.0.1 (Zhao et al., 2014). Further, pan-genome analysis of 11 *X. indica* strains and their pathogenic counterparts, *X. citri* pv. *citri* (*Xcc*) 306 and *X. oryzae* pv. *oryzae* (*Xoo*) BXO1, was performed using anvi’o platform v7.1 using DIAMOND to calculate gene similarity and MCL algorithm to identify gene clusters (Buchfink, Xie, & Huson, 2015; Eren et al., 2015; Van Dongen & Abreu-Goodger, 2012). The unique genes belonging to *X. indica* species and their pathogenic counterparts are functionally categorized in the Cluster of Orthologous Genes (COGs) using anvi’o integrated NCBI COGs database.

### Secretion systems and LPS O-antigen biosynthetic gene clusters

To check the presence/absence of secretion systems previously described in the genus *Xanthomonas*, their gene amino acid sequences were fetched from NCBI using locus tags. The gene sequences were taken as query to search their homologues in the genomes using tBLASTn v2.12.0+. The cut-offs for percent identity and query length coverage were set at >40% and >60%, respectively. The presence/absence heatmaps were created with TBtools v1.108 (Chen et al., 2020). To extract the lipopolysaccharide O-antigen biosynthetic gene cluster from the *X. indica* strains, *etfA* and *metB* gene sequences from a previous study were used as query (Patil & Sonti, 2004). *X. citri* pv. *citri* 306 (Xcc306) X-T4SS genes (XAC2612-XAC2623) were taken as query to search their homologues in the NPX genomes. Comparative gene clusters based on BLASTp were generated using Clinker v0.0.23 (Gilchrist & Chooi, 2021).

### Virulence assay and co-inoculation assay

The *X. indica* strains, PPL405 and PPL116 were inoculated on rice leaves to understand their pathogenicity status. Briefly, bacterial cultures were grown overnight in PS media (1% peptone and sucrose, pH 7.2) and were pelleted, washed with Milli-Q water, and adjusted to OD_600nm_ 1. Clip inoculation was performed on leaves of 60 to 80 days old susceptible rice cultivar Taichung Native 1 (TN1, a rice cultivar susceptible to bacterial blight disease). The NPX strains were also co-inoculated on rice leaves to understand if they protect against *Xoo* infection. Briefly, bacterial cultures were grown overnight in PS media (1% peptone and sucrose, pH 7.2), pelleted and washed with Milli-Q water. The NPX bacterial suspension equivalent for OD_600nm_ 1 measured at in 1ml was made for a volume of 0.5ml. This was mixed with 0.5 ml equivalent of wild-type Xoo strain BXO43 suspension just prior to clip inoculation. This ensures both NPX and Xoo are present to an OD_600nm_ 1 in the final mixture. As controls, rice leaves were clip inoculated with scissors dipped in either MilliQ water or OD_600nm_ 1 of BXO43 suspension. Leaves of 60 to 80 days old plants of rice cultivar Taichung Native 1 (TN1) were used for inoculation.

### Screening for antimicrobial gene clusters

BActeriocin Genome mining tooL (BAGEL4) was used to screen the NPX genome for the presence of bacteriocin-encoding gene clusters (van Heel et al., 2018). AntiSMASH v6.1.1 is used for identifying secondary metabolites such as ribosomally synthesized and post-translationally modified peptides (RiPPs) and nonribosomal peptide synthetases (NRPSs) (Blin et al., 2021).

## Results

### Genome sequencing, assembly, and annotation of *X. indica* isolates

Genomes of the type strain, PPL560^T^, and PPL568 were taken from the previous study from our lab (Rana et al., 2022). The type strains, *X. sontii* PPL1^T^, and *X. sacchari* CFBP4641^T^ were also sequenced in this study to obtain high-quality genomic data. Genomes for CFBP8445, *X. sacchari* F10, and LMG 8989 were obtained from the NCBI GenBank database. The draft genome size of strains ranges from 4.44 Mb to 5.04 Mb, with GC content ranging from 68.9% to 69.5%. The genomes have contig numbers between 1 to 191, with their genome coverage ranging from 125× to 1317×. All the genomes are 100% complete except for LMG 8989, which has 93% completeness. Detailed genome assembly and annotation statistics of all the genomes used in this study are given in **Table 1**.

**Table 1:**
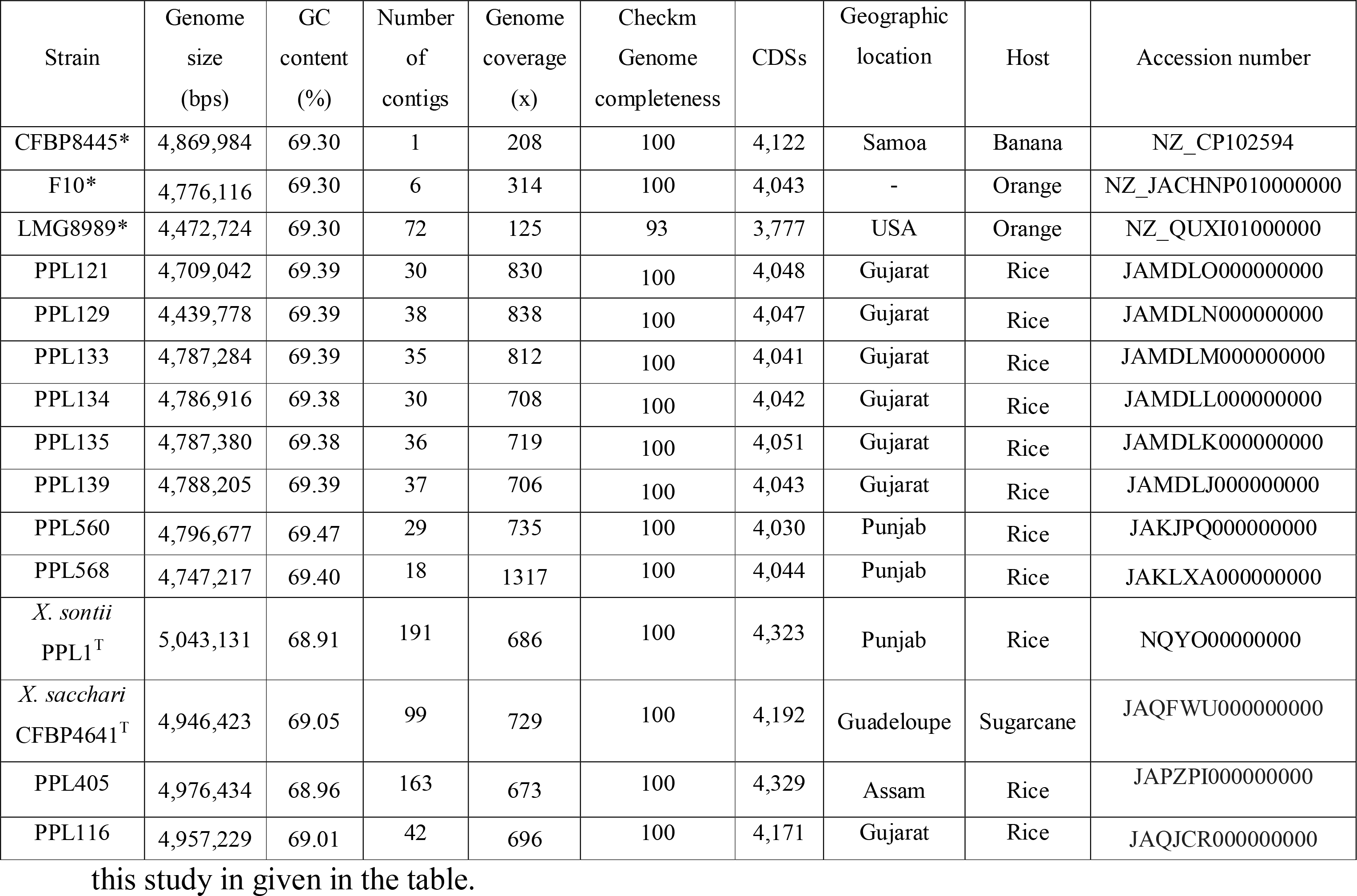
Genome assembly, annotation, and metadata statistics of the strains used in the study. ^*^Genome data obtained from the NCBI GenBank database. Genomic data for PPL560^T^ and PPL568 was taken from our previous study. Geographic location of Indian isolates from this study in given in the table.

### Phylo-taxonogenomic investigation into diversity *X. indica*

The phylo-taxonogenomic analysis revealed that CFBP8445 and LMG8989 strains isolated from banana and orange plants, respectively, also belong to *X. indica* (Bansal et al., 2020; Peduzzi et al., 2023). Further, *X. sacchari* F10 is also a misclassified *X. indica* strain isolated from an orange plant (**Figure 1, Table 2**). All the *X. indica* strains cluster in clade I group of the xanthomonads. Uncharacterised *Xanthomonas* strains D109 and NCPPB1132 belong to *X. sacchari* while SHU166, SHU308, SHU199, LMG12459, LMG12460, LMG12461, and LMG12462 strain belong to *X. sontii* (**Figure 1**). PPL405 isolated in this study belongs to *X. sacchari*, which is surprising given the fact that *X. sacchari* is known to infect sugarcane, whereas this strain was isolated from healthy rice seeds. Furthermore, all the *X. indica* strains, along with *X. sontii*, and *X. sacchari* and *X. albilineans*, the sugarcane pathogens, belong to clade 1B (Koebnik et al., 2021). Recently described novel species, *X. youngii* and *X. bonasiae* belong to the clade 1A along with *X. theicola, X. hycinthi, X. translucens* (Mafakheri, Taghavi, Zarei, Portier, et al., 2022). These two novel species are non-pathogenic on their host of isolation and lack type III secretion system (T3SS) and type III effectors (T3Es), however, they were able to induce hypersensitive response (HR) in the tobacco plant (Mafakheri, Taghavi, Zarei, Rahimi, et al., 2022). *X. maliensis* is another non-pathogenic species isolated from rice plant which belongs to the clade II group of xanthomonads (Triplett et al., 2015). Similarly, non-pathogenic strains reported from *X. arboricola* and *X. euroxanthea* are also clustered in the clade II group of xanthomonads along with their pathogenic counterparts (Cesbron et al., 2015; Martins et al., 2020). LMG9002 and LMG8992, non-pathogenic citrus isolates, were clustered in clade 1B in close proximity to *X. indica* strains. Isolates, GW, SS, and SI, clustered in clade 1A are also non-pathogenic in nature and isolated from ryegrass (Li et al., 2020). Here, the analysis reveals that NPXs are reported from both the major clades of xanthomonads. However, they are more frequently reported from the less explored clade I group.

**Table 2:**
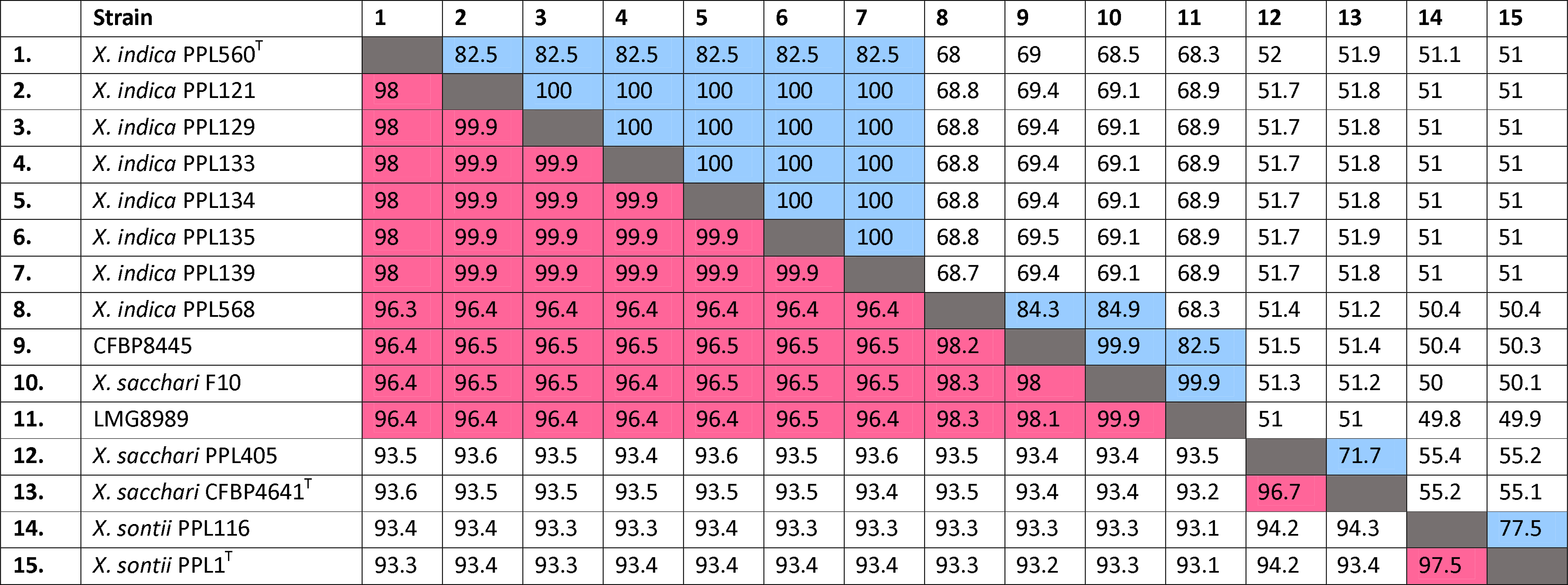
ANI and dDDH values of the *Xanthomonas* strains isolated from this study and from previous studies calculated amongst each other and with the type strains of *X. indica, X. sontii* and *X. sacchari*.

|  | Strain | 1 | 2 | 3 | 4 | 5 | 6 | 7 | 8 | 9 | 10 | 11 | 12 | 13 | 14 | 15 |
| --- | --- | --- | --- | --- | --- | --- | --- | --- | --- | --- | --- | --- | --- | --- | --- | --- |
| 1. | *X. indica* PPL560<sup>T</sup> |  | 82.5 | 82.5 | 82.5 | 82.5 | 82.5 | 82.5 | 68 | 69 | 68.5 | 68.3 | 52 | 51.9 | 51.1 | 51 |
| 2. | *X. indica* PPL121 | 98 |  | 100 | 100 | 100 | 100 | 100 | 68.8 | 69.4 | 69.1 | 68.9 | 51.7 | 51.8 | 51 | 51 |
| 3. | *X. indica* PPL129 | 98 | 99.9 |  | 100 | 100 | 100 | 100 | 68.8 | 69.4 | 69.1 | 68.9 | 51.7 | 51.8 | 51 | 51 |
| 4. | *X. indica* PPL133 | 98 | 99.9 | 99.9 |  | 100 | 100 | 100 | 68.8 | 69.4 | 69.1 | 68.9 | 51.7 | 51.8 | 51 | 51 |
| 5. | *X. indica* PPL134 | 98 | 99.9 | 99.9 | 99.9 |  | 100 | 100 | 68.8 | 69.4 | 69.1 | 68.9 | 51.7 | 51.8 | 51 | 51 |
| 6. | *X. indica* PPL135 | 98 | 99.9 | 99.9 | 99.9 | 99.9 |  | 100 | 68.8 | 69.5 | 69.1 | 68.9 | 51.7 | 51.9 | 51 | 51 |
| 7. | *X. indica* PPL139 | 98 | 99.9 | 99.9 | 99.9 | 99.9 | 99.9 |  | 68.7 | 69.4 | 69.1 | 68.9 | 51.7 | 51.8 | 51 | 51 |
| 8. | *X. indica* PPL568 | 96.3 | 96.4 | 96.4 | 96.4 | 96.4 | 96.4 | 96.4 |  | 84.3 | 84.9 | 68.3 | 51.4 | 51.2 | 50.4 | 50.4 |
| 9. | CFBP8445 | 96.4 | 96.5 | 96.5 | 96.5 | 96.5 | 96.5 | 96.5 | 98.2 |  | 99.9 | 82.5 | 51.5 | 51.4 | 50.4 | 50.3 |
| 10. | *X. sacchari* F10 | 96.4 | 96.5 | 96.5 | 96.4 | 96.5 | 96.5 | 96.5 | 98.3 | 98 |  | 99.9 | 51.3 | 51.2 | 50 | 50.1 |
| 11. | LMG8989 | 96.4 | 96.4 | 96.4 | 96.4 | 96.4 | 96.5 | 96.4 | 98.3 | 98.1 | 99.9 |  | 51 | 51 | 49.8 | 49.9 |
| 12. | *X. sacchari* PPL405 | 93.5 | 93.6 | 93.5 | 93.4 | 93.6 | 93.5 | 93.6 | 93.5 | 93.4 | 93.4 | 93.5 |  | 71.7 | 55.4 | 55.2 |
| 13. | *X. sacchari* CFBP4641<sup>T</sup> | 93.6 | 93.5 | 93.5 | 93.5 | 93.5 | 93.5 | 93.4 | 93.5 | 93.4 | 93.4 | 93.2 | 96.7 |  | 55.2 | 55.1 |
| 14. | *X. sontii* PPL116 | 93.4 | 93.4 | 93.3 | 93.3 | 93.4 | 93.3 | 93.3 | 93.3 | 93.3 | 93.3 | 93.1 | 94.2 | 94.3 |  | 77.5 |
| 15. | *X. sontii* PPL1<sup>T</sup> | 93.3 | 93.4 | 93.3 | 93.4 | 93.4 | 93.4 | 93.4 | 93.3 | 93.2 | 93.3 | 93.1 | 94.2 | 93.4 | 97.5 |  |

**Figure 1.**
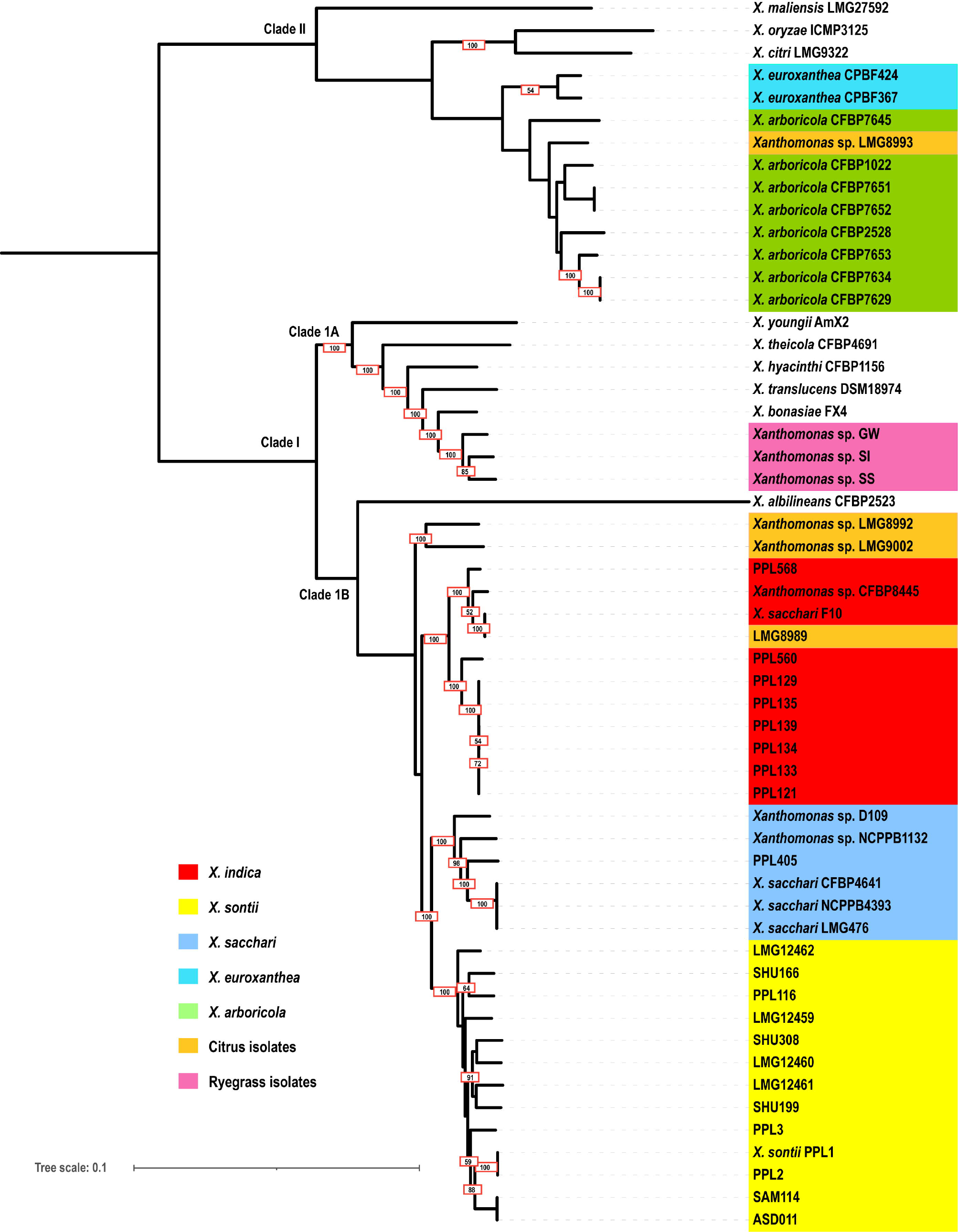
Mid-rooted core gene phylogeny of NPX isolates from this study with other NPX strains and some important representatives of the genus *Xanthomonas*. The strains are clustered in two clades, clade I and clade II. Clade I further have two sub-clades, clade IA and clade IB. Bootstraps values are highlighted on the branches. The *Xanthomonas* species with NPX strains are marked with different colors.

All the *X. indica* strains shared 96.4% to 99.9% ANI values between them. Six of the isolates, PPL121, PPL129, PPL133, PPL134, PPL135, and PPL139, shared approximately 99.9% ANI values with each other. The ANI values and SNP phylogeny revealed the clonal nature of PPL121, PPL129, PPL133, PPL134, PPL135, and PPL139, which were clustered together. The SNP distance matrix created with CSI phylogeny shows the number of SNP differences amongst PPL121, PPL129, PPL133, PPL134, PPL135, and PPL139 ranges from 4 to 22 (**Supplementary figure 1a, 1b)**. PPL116 belongs to *X. sontii* based on the ANI value, which was 97.5% with *X. sontii* PPL1^T^. PPL405 shares a 96.7% ANI value with *X. sacchari* CFBP4641^T^. The dDDH values of *X. indica* isolates with *X. indica* PPL560^T^ are in the range of 68.3% to 82.5%. For *X. sacchari* F10, CBP8445, and LMG8989, the dDDH values are slightly below the cut-off (≥70%), however, based on their phylogenetic analysis and their ANI values, these strains belong to *X. indica* (**Table 2**). PPL121, PPL129, PPL133, PPL134, PPL135, and PPL139 share 100% dDDH values with each other. LMG8989 and F10 share dDDH and ANI values of 99.9% and 98% with each other, respectively, which is interesting as both are citrus isolates and are forming a monophyletic subclade in the *X. indica* clade. All the ANI and dDDH values of the rice isolates from this study and their closest *Xanthomonas* strains are given in **Table 2**.

### Comparative genomics of *X. indica* isolates reveal open pangenome and genome adaptation

Pan-genome analysis of *X. indica* revealed the pan-genome size to be 5,023, with 3,222 core genes shared by all 11 genomes. The species has an open pan-genome, where the profile curve expands with the addition of genomes, the pan-genome keeps increasing, and core gene content remains stable with a slight decrease (**Figure 2a)**. *X. indica* strains are diverse with a significant number of unique genes. PPL560 has 144, PPL568 has 209, and F10 and CFBP8445 each has 12 unique genes. Further, due to their clonal nature, PPL121, PPL129, PPL133, PPL134, PPL135, and PPL139 have the least number of unique genes (**Figure 2b). Figure 2c** shows the pan-genome analysis of 11 *X. indica* strains with *X. citri* pv. *citri* (Xcc 306) and *X. oryzae* pv. *oryzae* (Xoo BXO1). The pan-genome size of 13 genomes is 6,765 gene clusters, and they share 2,025 core gene clusters. The non-pathogenic *X. indica* strains and pathogenic strains (Xoo BXO1 and Xcc 306) have 820 and 507 unique gene clusters, respectively. Further, the unique genes of pathogenic and non-pathogenic strains are categorized into COG categories **(Figure 2d)**. The NPX strains have a greater number of unique genes in the defense mechanisms (V), Secondary metabolites biosynthesis, transport and catabolism (M), and Extracellular structures (W), whereas the pathogenic strains carry a greater number of unique genes in the Intracellular trafficking, secretion, and vesicular transport (U) categorizes. Unique genes categorized into Mobilome: prophages, transposons (X) were exclusively present in the pathogenic strains **(Figure 2d)**.

**Figure 2.**
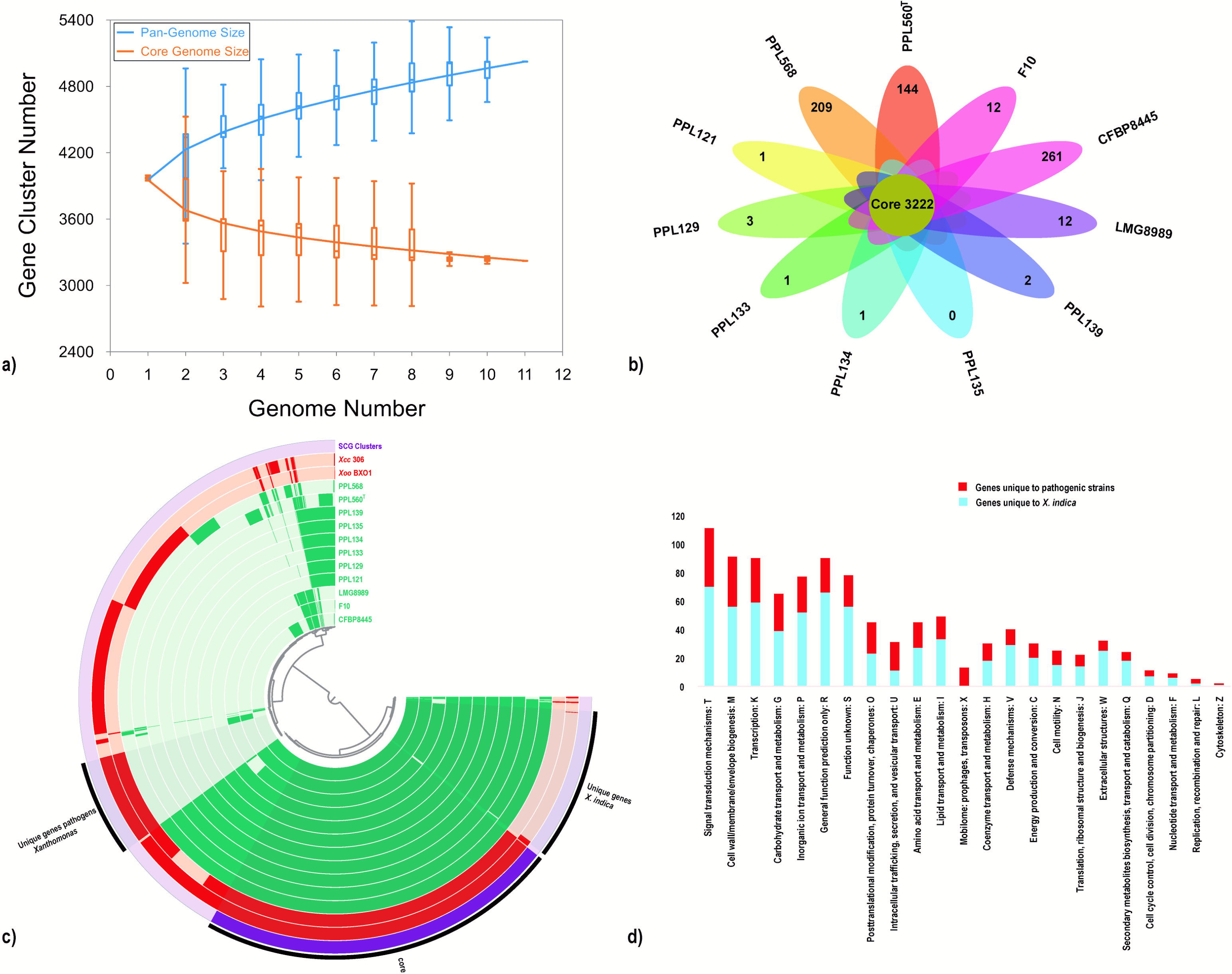
Pan-genome analysis of *X. indica* strains. **a)** Pangenome profile curve with x-axis and y-axis representing genome number and gene cluster numbers, respectively. The colored boxes depict pan-genome size (blue) and core-genome size (orange). **b)** The flower plot shows number of core genes in the centre and the flower petals have number of unique genes of each strain. **c)** Pan-genome profile of *X. indica* strains with their pathogenic counterparts, i.e. *Xoo* and *Xcc*. Each circular ring represents each genome. Green and red colored rings represent *X. indica* strains and pathogenic strains, respectively. Core genes, unique genes to *X. indica*, and pathogenic strains are highlighted with black colored bars. **d)** Functional categorization of genes unique to *X. indica* strains and pathogenic strains. X-axis and y-axis represent COG categories and number of genes, respectively. Color coding scheme is given in the figure.

### Protein secretion systems and virulence loci in *X. indica*

Xanthomonads are known to carry six secretion systems that function in promoting attachment, virulence, and interbacterial competition. Type I secretion system (T1SS) associated RaxA, RaxST, and RaxX were absent, while RaxB and RaxC, along with a sulfotransferase-like protein and a two-component regulatory system consisting of RaxH and RaxR genes of Xoo were present in the *X. indica* strains. (Burdman, Shen, Lee, Xue, & Ronald, 2004). T1SS genes of *Xcc* were absent in the *X. indica* strains except for PctB, a homologue of RaxB (Luu et al., 2019) **(Figure 3a)**. As seen in Xoo, *X. indica* strains have *xps* type II secretion system (T2SS) and lack *xcs* T2SS (Lu et al., 2008; Moreira et al., 2004; Szczesny et al., 2010). The strains also carry XcsE gene of *xcs* T2SS at very low identity values **(Figure 3b)**. The *xps* T2SS releases cell wall degrading enzymes (CWDEs) such as lipases/esterase, cellulases, proteases, xylanases, endoglucanases, pectate lyase, and polygalacturonases (Jha, Rajeshwari, & Sonti, 2005; Potnis et al., 2011; Solé et al., 2015). X. indica strains carry xylanases (Xyn51A and Xyn5A), cellulase (ClsA), endoglucanase (Egl), pectate lyases (Pel, Pel4A), pectin methylesterase (PemB), glucosidase (CelD), protease (ClpA), and glucuronidase (aguA) **(Figure 4g)**. T3SS and T3Es that were absent in the type strain are also absent in all the *X. indica* strains as mentioned in the previous study (Rana et al., 2022) **(Figure 3c, Figure 3h)**. Interestingly, *X. indica* strains carry homologues of the T4SS from *Xcc*, but this gene cluster is absent from Xoo (**Figure 3d)**. T5SS mainly play in promoting adhesion of *Xanthomonas* strains and thus support initial interaction with the hosts. *X. indica* strains carry EstA (lipase/esterase), a monomeric autotransporter, and FhaB/FhaC genes, PPL121-PPL139 strains lack FhaB (Gottig, Garavaglia, Garofalo, Orellano, & Ottado, 2009; Guérin, Bigot, Schneider, Buchanan, & Jacob-Dubuisson, 2017; Meuskens, Saragliadis, Leo, & Linke, 2019). Type V secretion system (T5SS) adhesins (Xad and YapH) were absent in the *X. indica* strains, with the exception of XadB, which is present in PPL568, LMG8989, F10, CFBP8445 **(Figure 3e)**. The xanthomonads are known to carry type VI secretion systems (T6SS), T6SS^i3*^, T6SS^i3***^, and T6SS^i4^ types of T6SSs, *X. indica* strains lack all three of them (Bayer-Santos, Ceseti, Farah, & Alvarez-Martinez, 2019) (**Figure 3f)**. The primary regulator proteins, HrpX and HrpG, which are directly or indirectly involved with the other T3SS-associated regulator proteins, are absent in *X. indica* strains (Alvarez-Martinez et al., 2021; Büttner & Bonas, 2010). All the *X. indica* strains carry two-component regulatory systems, ColRS, PhoPQ, PhoBR (S.-W. Lee et al., 2008; Zheng et al., 2018). Further, a diffusible signal factor (DSF) based regulatory system consisting of Rpf proteins, RpfF, RpfB, and RpfCG is also present in *X. indica* species (Andrade et al., 2006; Büttner & Bonas, 2010). **Figure 3i** depicts the presence/absence profile of some important regulatory functions in *X. indica*.

**Figure 3.**
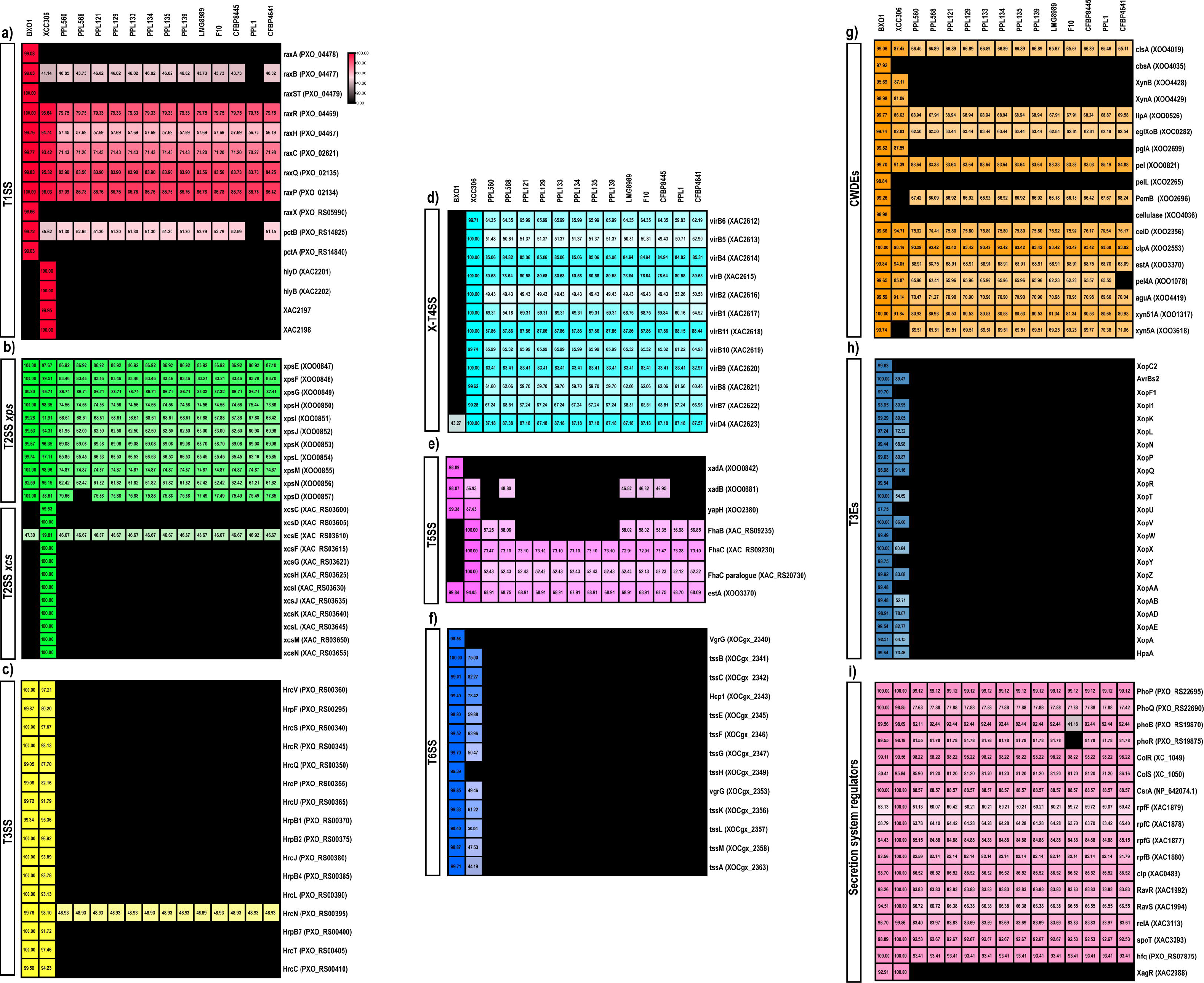
The presence/absence profile of secretion systems and their regulators. **a)** T1SS. **b)** *xps* and *xcs* T2SS. **c)** T3SS. **d)** T4SS. **e)** T5SS. **f)** T6SS. **g)** CWDEs. **h)** T3Es. **i)** Regulators. The black colored boxes depict the absence of genes. The color gradients represent identity values, where the gene identity values are given in the boxes. NCBI locus tags of the genes are given in brackets against gene names.

### Species-level variation in the LPS O-antigen biosynthetic gene cluster

The Lipopolysaccharide (LPS) constitutes an important part of the outer membrane in gram-negative bacteria that is recognized by plants as a Pathogen-Associated Molecular Pattern (PAMP) (Raetz & Whitfield, 2002). In the case of plant pathogens, they are involved in eliciting immune response in plants, and are important virulence factors (Dow, Osbourn, Wilson, & Daniels, 1995; Girija et al., 2017). Previous studies have reported their hypervariability at the species level in pathogenic Xanthomonads (Singh, Bansal, Kumar, & Patil, 2022; Wasukira et al., 2014). *X. indica* species was found to carry three different kinds of LPS gene clusters **(Figure 4)**. PPL568, F10, LMG898, and PPL121 have BXO8 type of LPS cassette, however, two hypothetical protein, class I SAM-dependent methyltransferase, and an IS elements (IS3) present in the BXO8 LPS cassette is absent from *X. indica* strains (Singh et al., 2022). Further, PPL560 and CFBP8445 have two different kinds of LPS cassettes from BXO8. The LPS cassette from PPL560 is similar to the XVV type LPS cassette present in *X. sacchari* CFBP4641 on the *metB* side of the LPS cassette, PPL560 has 3 glycosyl transferases, *gtrA*, and a hypothetical protein-encoding genes on *etfA* side which do not share similarity to any other LPS locus (Singh et al., 2022) **(Figure 4)**. CFBP8445 has an LPS cassette that shows similarity to both BXO8 and XOC type of LPS cassettes, however, this LPS locus is reduced in size when compared to the other two and has a hypothetical gene and a FkbM methyltransferase that does not show similarity to any other gene from LPS the gene cluster present in *Xanthomonas*. Detailed LPS gene cluster annotation and their GC% content are given in **Supplementary table 1**.

**Figure 4.**
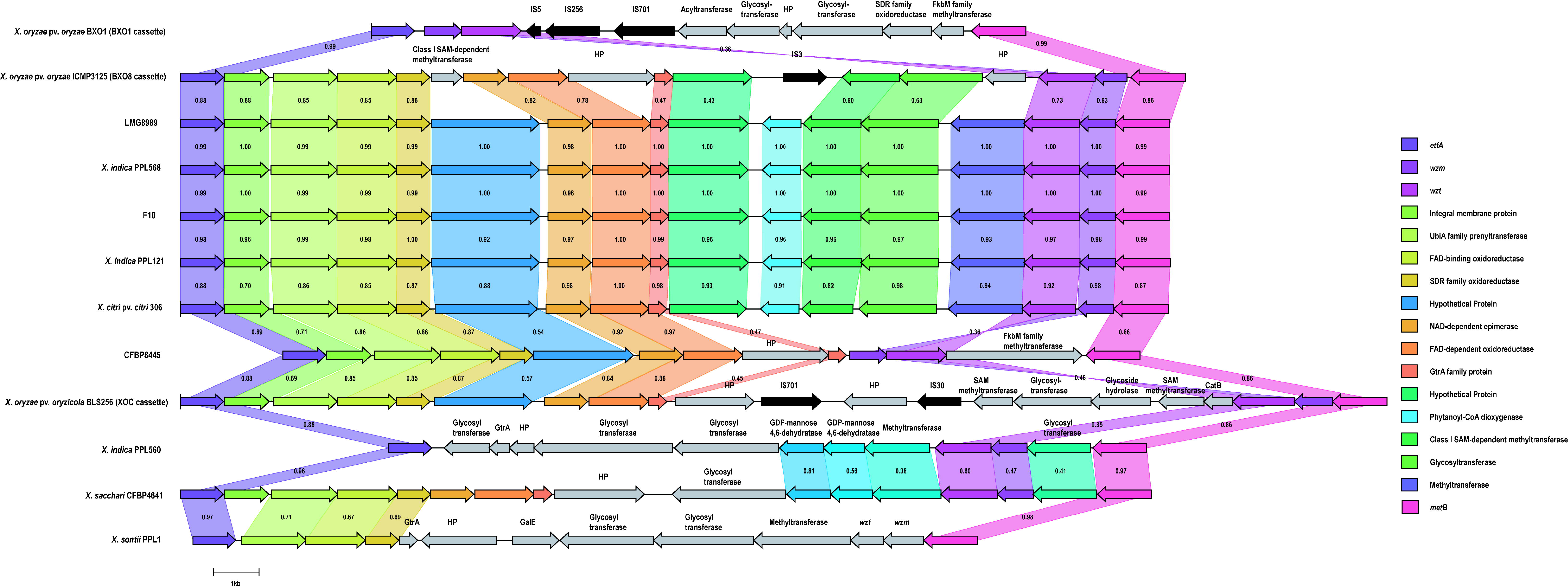
Comparison between *X. indica* isolates and previously reported LPS gene clusters. Gene homologues across the strains are represented with same colored arrows. Protein homology is shown by connecting regions along with identity values. Gene annotations are given on the right-hand side of the image. Gene unique to a particular strain are annotated on the arrows.

### Diverse *X. indica* strains provide protection to rice plants against leaf blight pathogen

Since all the *X. indica* strains lacked T3SS, we assume these strains would also be non-pathogenic in nature. To know the pathogenicity status of the newly isolated strains, they were clip inoculated on rice plants. As expected, none of the isolates generated virulence lesions compared to the virulence-proficient Xoo strain that was used as a control **(Figure 5a, 5b)**.

**Figure 5.**
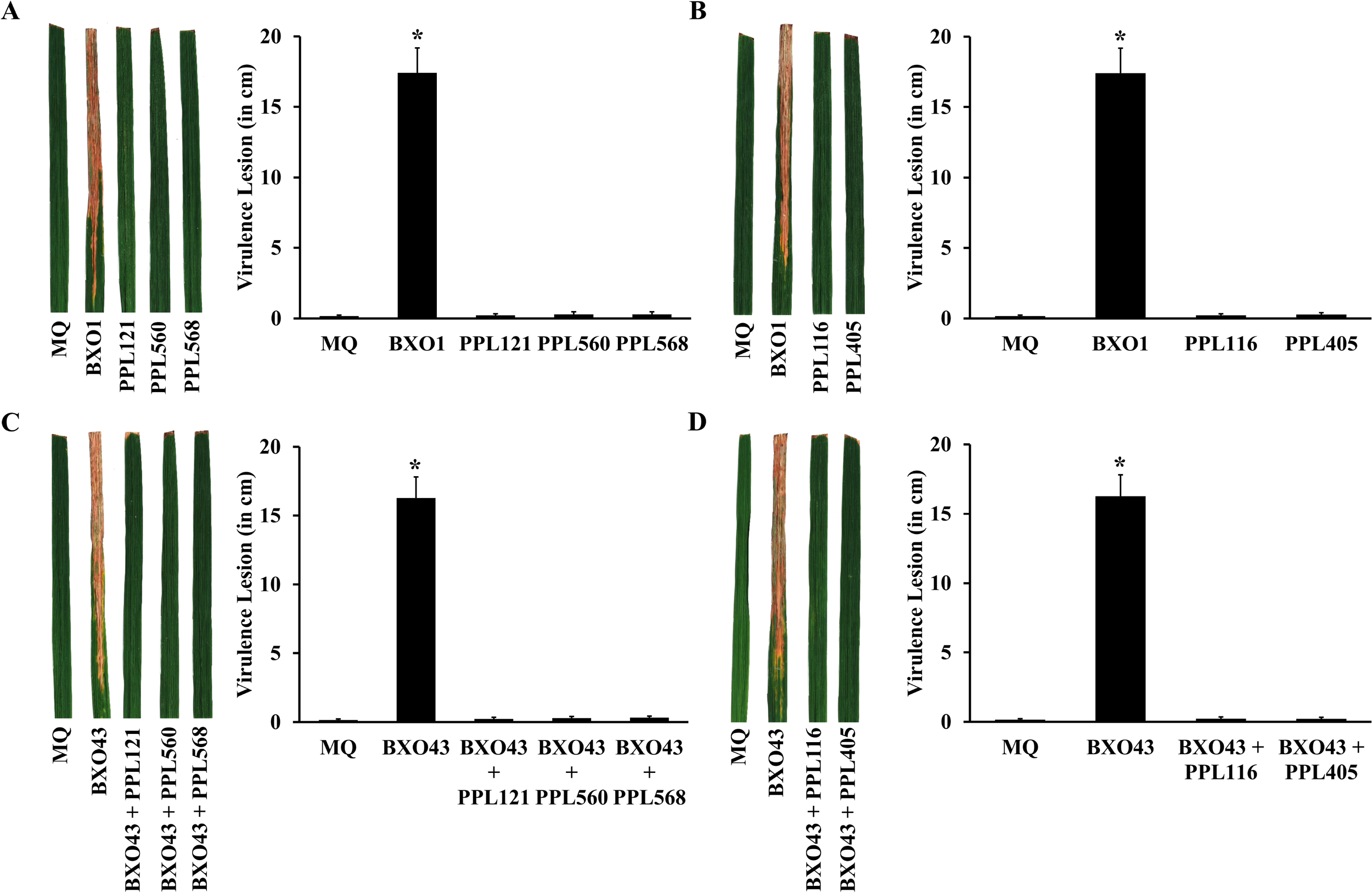
Checking pathogenicity and possibility of biocontrol nature of NPX strains. The bacterial strains were inoculated as described in the materials and methods. Sample composite image of infected rice leaves and bar graph of average virulence lesion lengths observed for **(A)** *X. indica* strains, PPL121, PPL560, PPL568 and **(B)** *X. sontii* strain PPL116 and *X. sacchari* strain PPL405 compared with wild type Xoo, BXO1. Sample composite image of infected rice leaves and bar graph of average virulence lesion lengths observed for co-inoculation of wild type Xoo strain BXO43 with **(C)** *X. indica strains*, PPL121, PPL560, PPL568 and **(D)** *X. sontii* strain PPL116 and *X. sacchari* strain PPL405. Error bars used are standard deviations calculated from a minimum of 14 leaves. The * labelling of the column indicates a significant difference in lesion length using unpaired two-tailed student’s *t*-test (*p*-value < 0.01).

**Figure 6.**
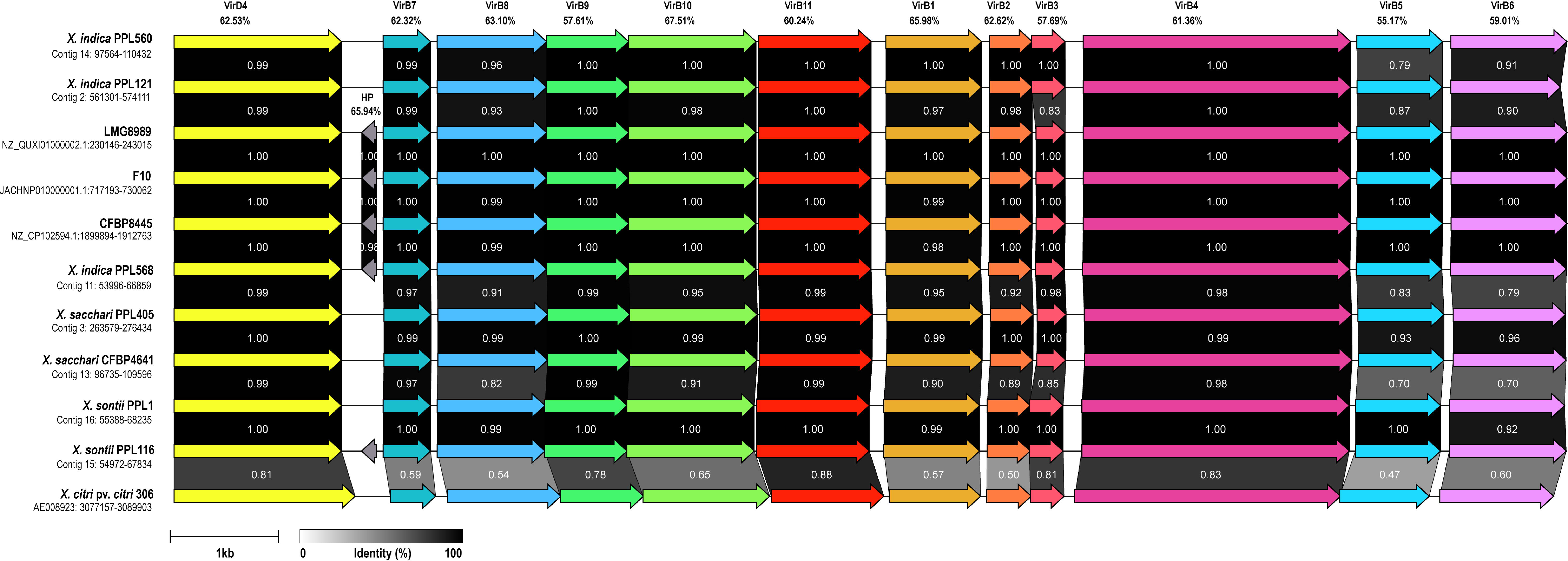
X-T4SS encoding gene cluster. Gene homologues across the strains are represented with same colored arrows. Connecting regions have amino acid identity values. Gene annotations are given on the top of the arrows along with their GC% content.

Since NPX strains seem to be co-evolved alongside pathogenic strains, we asked the question if they are in any manner beneficial to the host plant. One such possibility is that NPX strains provide protection against the virulence-proficient pathogenic counterparts. To understand if any of these NPX strains affected virulence of Xoo, we co-inoculated suspensions of NPX strains and *Xoo* wild-type strain BXO43. Interestingly, no virulence lesions were observed in the leaves that were co-inoculated **(Figure 5c, 5d)**. This suggests that there is a possibility that NPX strains may inhibit the growth of Xoo its *in-planta*. Importantly, we had earlier also observed a bio-protection property in *X. sontii* and *X. sacchari*.

### Putative genomic loci associated with protection against Xoo infection

Only one of the *X. indica* strains, PPL560 harbours a gene cluster encoding for Lanthipeptide class IV indicating another locus in plant protection activity. Similarly, several strains of *X. indica* strains have a 20 kb gene cluster encoding for sactipeptides, which is absent in related *X. sontii* and *X. sacchari* isolates. AntiSMASH analysis revealed the presence of a common gene cluster of 10.8 kb belonging to RiPP-like class **(Supplementary table 3)**. Strains belonging to *X. sontii* and *X. sacchari* have multiple NRPS encoding genes, however *X. indica* strains do not have any NRPS gene cluster. Interestingly, all the strains of *X. indica* along with *X. sontii*, and *X. sacchari* type strain, have a 20.3 kb gene cluster of ZoocinA class, which encodes for M23 family metallopeptidase **(Supplementary table 2)**.

T4SS was reported to be involved in contact-based bacterial killing in the citrus pathogen, *Xanthomonas citri*, additionally *Stenotrophomonas maltophilia*, an opportunistic human pathogen is also known to use T4SS for bacterial antagonism (Bayer-Santos, Cenens, et al., 2019; Souza et al., 2015). T4SS is made up of 12 subunits, VirB1-VirB11 and VirD4 which form a transmembrane multimodular complex along with extracellular pili (Sgro et al., 2019). The chromosomally associated T4SS (X-T4SS) helps in bacterial antagonism with the transfer of toxins into competitor strains (Sgro et al., 2019; Souza et al., 2015). **Figure 3d** depicts the presence/absence of the X-T4SS homologues of Xcc 306 in the *X. indica, X. sontii*, and *X. sacchari* strains from the study along with their percent identity values. Based on the cut-offs the X-T4SS is present in all the plant-tested strains for the bio-protection potential. Furthermore, the X-T4SS is present in the *X. indica* strains from other studies, such as LMG8989, F10, and CFBP8445, as well as the type strains of *X. sontii* and *X. sacchari*. The X-T4SS gene cluster of representatives of *X. sontii, X. sacchari*, and *X. indica* with *Xcc* 306 is shown in **Figure 5**. These three *Xanthomonas* species have all the components of X-T4SS with *virB* locus (*virB1*-*virB11*) and *virD4*. The X-T4SS gene cluster lies in the contig 14 of PPL560^T^ at coordinates, 97,564-110,432, contig 16 of PPL1^T^ at coordinates, 55,388-68,235, and contig 3 of PPL405 from coordinates 263,579 to 276,434. Interestingly, the X-T4SS is absent from Xoo strain BXO1 (**Figure 5**), BXO43 which was used for the co-inoculation experiment is a rif^r^ derivative of BXO1 (Ray, Rajeshwari, & Sonti, 2000). The presence of X-T4SS in the NPX and its absence in the Xoo gives a selective antagonistic advantage to NPX strains over *Xoo*.

## Discussion

Similar to the complex world of phytopathogenic *Xanthomonas*, it is becoming increasingly clear that there is a world of non-pathogenic *Xanthomonas* (NPX) species (Bansal et al., 2021; Bansal et al., 2020; Rana et al., 2022). Although these NPX strains belong to the same genus, they differ greatly from their pathogenic counterparts, as they lack major virulence factors and might also be involved in commensal relationships with the host plants (Mafakheri, Taghavi, Zarei, Rahimi, et al., 2022; Rana et al., 2022). In this context, rice is a good model plant as only one species, i.e. *X. oryzae* was known for the last 100 years, but with the beginning of this century, three non-pathogenic species have been reported indicating the importance of NPX species (Bansal et al., 2021; Rana et al., 2022; Triplett et al., 2015). Unlike pathogenic species, individual or species studies to understand strain or population level genomic diversity in NPX community is lacking and there is a need for more focussed studies. Being a model and staple food crop, it is important to study the diversity and functional role of NPX species or community from rice plants. Our study fills this gap with isolation, sequencing, and genomic characterization of multiple strains of the newest NPX species from the rice *Xanthomonas* community. Interestingly, the limited number of genome sequences revealed significant and extensive diversity suggesting the importance of sequencing strains from diverse geographic regions. The inclusion of the lineage consisting of *X. indica* strains not only provide clarity regarding to get phylogeny of the two major clades I and II found in the genus *Xanthomonas* but also confirmed the existence of and relationship amongst sub-clades in Clade-I consisting of other NPX species along with its closest relatives.

The *X. indica* strains lack major pathogenicity factors such as T3SS, T3Es, T6SS, adhesins and some important CWDEs supporting a distinct non-pathogenic lifestyle of NPX species. The presence of several NPX species with substantial levels of diversity suggests that these are important members of the rice microbiome with a potentially important role in host health. In this context, it is pertinent to note that while *X. sontii* strains were reported primarily from rice, *X. indica* strains have also been reported from other plants like citrus and banana, making it an important NPX species. This also suggests the existence of autochthonous and allochthonous species of NPX. The open pangenome and the presence of three different kinds of LPS gene clusters in *X. indica* species suggest ongoing functional diversification. The finding of these diverse *X. indica* strains protecting rice from a pathogenic *Xanthomonas* strains points to its role in co-adaptation with plants. NRPS are also well-known antimicrobial biosynthetic genomic loci, and the presence of NRPS cassette(s) in some of the NPX is indicative of their importance. However, the absence of a common NRPS locus that can be associated with bio-protection properties indicates that they may play other important roles. Interestingly, the conservation of this function and the association of a bacterial-killing type IV protein secretion system in diverse strains of *X. indica* and its relatives from a major clade of Xanthomonads including *X. sontii*, another NPX species of rice, indicate evolving lifestyle and adaptation in these group of bacteria. Homologues of X-T4SS are also found in other xanthomonads, such as *X. citri* pathovars, *X. campestris* pathovars, *X. albilineans* and *X. arboricola* strains. Moreover, a similar *virB* cluster is also present in some other genera of the order Xanthomonadales, *Stenotrophomonas, Pseudoxanthomonas, Luteimonas, Lysobacter*, and *Luteibacter* (Sgro et al., 2019). Further, the strains with the bio-protection potential also carry a common RiPP-like gene cluster and a M23 family metallopeptidase, which are also reported to be associated with anti-bacterial properties (Abramov et al., 2022; Heilbronner, Krismer, Brötz-Oesterhelt, & Peschel, 2021). Further studies of these loci using genetic approaches and expression studies are underway.

## Authors’ Contributions

RR isolated the strains, performed identification, and genome analysis. VNM performed plant virulence and co-inoculation assay. RR drafted the manuscript with inputs from VNM, RVS, HKP and PBP. PBP planned and participated in the design of the study along with RR, HKP, and RVS. PBP, HKP, and RVS applied for funding and coordinated the study. All authors read and approved the manuscript.

