## Supplementary data for "Comparative genomics-based insights into diversification and bio-protection function of *Xanthomonas indica*, a non-pathogenic species of rice"

**Supplementary figure 1:** Single nucleotide polymorphism (SNP) analysis using by CSI phylogeny v1.4. a) SNP phylogeny, highlighted strains are clones of each other b) SNP matrix, strain names are highlighted in yellow colour. Blue coloured boxes show least SNPs differences across the *X. indica* stains.

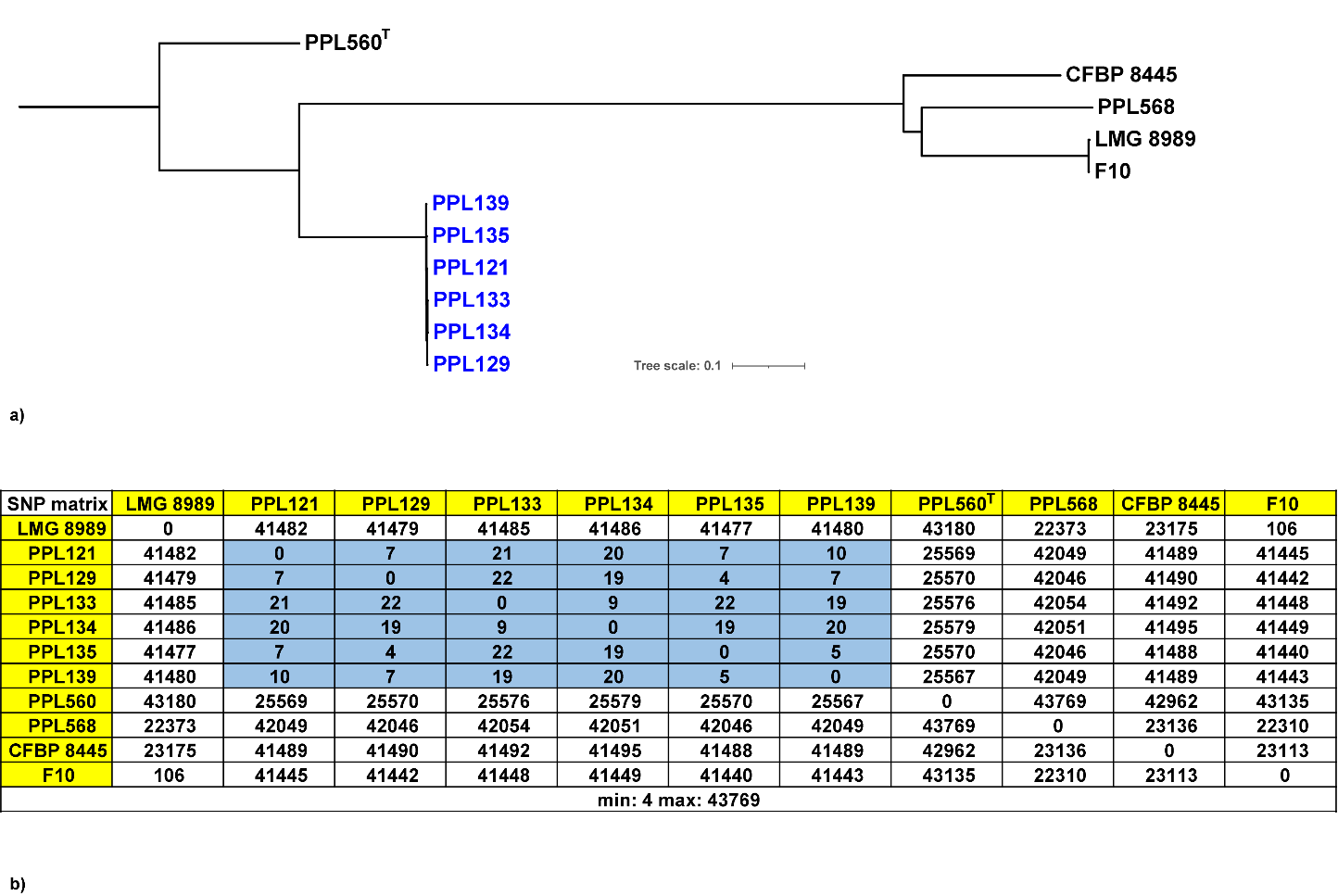

**Supplementary Table 1:** The LPS cassettes from PPL560, PPL568, and CFBP8445 with the locus tag, GC content and GenBank annotation of each gene in the cassette.

| **PPL568 LPS cassette** |  |  |
| --- | --- | --- |
| **Locus tag** | **GC content (%)** | **GenBank Annotation** |
| L3067_RS10280 | 70.48 | electron transfer flavoprotein subunit alpha/FixB family protein |
| L3067_RS10285 | 72.9 | lysylphosphatidylglycerol synthase transmembrane domain-containing protein |
| L3067_RS10290 | 67.9 | UbiA family prenyltransferase |
| L3067_RS10295 | 71.55 | FAD-binding oxidoreductase |
| L3067_RS10300 | 70.91 | SDR family oxidoreductase |
| L3067_RS10305 | 62.02 | hypothetical protein |
| L3067_RS10310 | 61.26 | NAD-dependent epimerase/dehydratase family protein |
| L3067_RS10315 | 62.16 | NAD(P)/FAD-dependent oxidoreductase |
| L3067_RS10320 | 51.88 | GtrA family protein |
| L3067_RS10325 | 59.92 | hypothetical protein |
| L3067_RS10330 | 55.98 | phytanoyl-CoA dioxygenase family protein |
| L3067_RS10335 | 57.91 | hypothetical protein |
| L3067_RS10340 | 60.43 | glycosyltransferase family 2 protein |
| L3067_RS10345 | 58.02 | class I SAM-dependent methyltransferase |
| L3067_RS10350 | 55.64 | ABC transporter ATP-binding protein |
| L3067_RS10355 | 55.81 | ABC transporter permease |
| L3067_RS10360 | 69.59 | cystathionine gamma-synthase |
| **PPL560 LPS cassette** |  |  |
| **Locus tag** | **GC content (%)** | **GenBank Annotation** |
| L3V74_RS08945 | 69.02 | cystathionine gamma-synthase |
| L3V74_RS08950 | 59.28 | glycosyltransferase family 4 protein |
| L3V74_RS08955 | 54.02 | ABC transporter permease |
| L3V74_RS08960 | 56.61 | ABC transporter ATP-binding protein |
| L3V74_RS08965 | 60.18 | methyltransferase domain-containing protein |
| L3V74_RS08970 | 60.96 | GDP-mannose 4,6-dehydratase |
| L3V74_RS08975 | 55.34 | GDP-mannose 4,6-dehydratase |
| L3V74_RS08980 | 57.16 | glycosyltransferase family 39 protein |
| L3V74_RS08985 | 54.67 | glycosyltransferase |
| L3V74_RS08990 | 61.72 | hypothetical protein |
| L3V74_RS08995 | 59.06 | GtrA family protein |
| L3V74_RS09000 | 55.92 | glycosyltransferase family 2 protein |
| L3V74_RS09005 | 69.42 | electron transfer flavoprotein subunit alpha/FixB family protein |
| **CFBP8445 LPS cassette** |  |  |
| **Locus tag** | **GC content (%)** | **GenBank Annotation** |
| NUG21_RS17605 | 70.38 | electron transfer flavoprotein subunit alpha/FixB family protein |
| NUG21_RS17610 | 73 | lysylphosphatidylglycerol synthase transmembrane domain-containing protein |
| NUG21_RS17615 | 67.45 | UbiA family prenyltransferase |
| NUG21_RS17620 | 71.4 | FAD-binding oxidoreductase |
| NUG21_RS17625 | 71.33 | SDR family oxidoreductase |
| NUG21_RS17630 | 55.57 | hypothetical protein |
| NUG21_RS17635 | 63.17 | NAD-dependent epimerase/dehydratase family protein |
| NUG21_RS17640 | 62.03 | NAD(P)/FAD-dependent oxidoreductase |
| NUG21_RS17645 | 61.84 | hypothetical protein |
| NUG21_RS17650 | 59.7 | GtrA family protein |
| NUG21_RS17655 | 57.26 | ABC transporter permease |
| NUG21_RS17660 | 59.9 | ABC transporter ATP-binding protein |
| NUG21_RS17665 | 61.45 | FkbM family methyltransferase |
| NUG21_RS17670 | 69.7 | cystathionine gamma-synthase |

**Supplementary table 2:** BAGEL4 analysis of *Xanthomonas* strains sequenced in this study showing presence of bacteriocin encoding gene clusters, their class, genomic region, and size. A gene cluster common to the strains tested for plant bio-protection assay is highlighted (orange colour).

| **Strain** | **Class** | **Genomic region (Contig: start-end)** | **Size (kb)** |
| --- | --- | --- | --- |
| **PPL560** | Zoocin A (M23 family metallopeptidase) | Contig 7:45759-66065 | 20.3 |
| **PPL560** | Sactipeptides | Contig 9:190451-210451 | 20 |
| **PPL560** | Lanthipeptide class IV | Contig 15:128564-148564 | 20 |
| **PPL568** | Zoocin A (M23 family metallopeptidase) | Contig 2:340484-360790 | 20.3 |
| **PPL568** | Sactipeptides | Contig 10:165902-185902 | 20 |
| **PPL121** | Zoocin A (M23 family metallopeptidase) | Contig 4:75153-95459 | 20.3 |
| **PPL121** | Sactipeptides | Contig 7:188792-208792 | 20 |
| **PPL129** | Sactipeptides | Contig 7:12224-32224 | 20 |
| **PPL129** | Zoocin A (M23 family metallopeptidase) | Contig 10:71618-91924 | 20.3 |
| **PPL133** | Zoocin A (M23 family metallopeptidase) | Contig 6:71618-91924 | 20.3 |
| **PPL133** | Sactipeptides | Contig 8:12224-32224 | 20 |
| **PPL134** | Zoocin A (M23 family metallopeptidase) | Contig 1:860279-880585 | 20.3 |
| **PPL134** | Sactipeptides | Contig 5:188792-208792 | 20 |
| **PPL135** | Sactipeptides | Contig 7:188792-208792 | 20 |
| **PPL135** | Zoocin A (M23 family metallopeptidase) | Contig 10:75152-95458 | 20.3 |
| **PPL139** | Sactipeptides | Contig 6:12224-32224 | 20 |
| **PPL139** | Zoocin A (M23 family metallopeptidase) | Contig 10:75152-95458 | 20.3 |
| **F10** | Sactipeptides | JACHNP010000001.1:527000-547000 | 20 |
| **F10** | Zoocin A (M23 family metallopeptidase) | JACHNP010000001.1:1702583-1722889 | 20.3 |
| **LMG8989** | Sactipeptides | NZ_QUXI01000003.1:164705-184705 | 20 |
| **LMG8989** | Zoocin A (M23 family metallopeptidase) | NZ_QUXI01000011.1:467-20773 | 20.3 |
| **CFBP8445** | Zoocin A (M23 family metallopeptidase) | NZ_CP102594.1:2934980-2955286 | 20.3 |
| **PPL1** | Zoocin A (M23 family metallopeptidase) | Contig 7:44630-64936 | 20.3 |
| **PPL116** | Zoocin A (M23 family metallopeptidase) | Contig 3:342526-357314 | 14.8 |
| **CFBP4641** | Zoocin A (M23 family metallopeptidase) | Contig 19:90950-105650 | 14.7 |
| **PPL405** | Zoocin A (M23 family metallopeptidase) | Contig 12:11837-32143 | 20.3 |

**Supplementary Table 3:** AntiSMASH v6.1.1 analysis of Xanthomonads from this to show the presence of secondary metabolites, their type, genomic region, and size. A gene cluster common to the strains tested for plant bio-protection assay is highlighted (orange colour).

| **Strain** | **Type** | **Genomic region (Locus tag, start: end)** | **Size (kb)** |
| --- | --- | --- | --- |
| **PPL560** | RiPP-like | L3V74_06815: L3V74_06865 | 10.8 |
| **PPL560** | Lanhipeptide class IV | L3V74_16960: L3V74_16880 | 22.7 |
| **PPL568** | RiPP-like | L3067_02925: L3067_02970 | 10.8 |
| **PPL121** | RiPP-like | M3U22_00765: M3U22_00815 | 10.8 |
| **PPL129** | RiPP-like | M3U23_09875: M3U23_09925 | 10.8 |
| **PPL133** | RiPP-like | M3S05_02170: M3S05_02220 | 10.8 |
| **PPL134** | RiPP-like | M3U25_00765: M3U25_00815 | 10.8 |
| **PPL135** | RiPP-like | M3U24_09370: M3U24_09420 | 10.8 |
| **PPL139** | RiPP-like | M3S04_00765: M3S04_00815 | 10.8 |
| **F10** | RiPP-like | FHR56_RS16630: FHR56_RS16680 | 10.8 |
| **LMG8989** | RiPP-like | DYQ91_RS03190: DYQ91_RS03240 | 10.8 |
| **CFBP8445** | RiPP-like | NUG21_RS07495: NUG21_RS07540 | 9.8 |
| **CFBP8445** | RiPP-like | NUG21_RS14585: NUG21_RS14610 | 10.9 |
| **PPL1** | NRPS | CJ027_004295: CJ027_004375 | 30.9 |
| **PPL1** | NRPS | CJ027_008040: CJ027_008120 | 22.8 |
| **PPL1** | RiPP-like | CJ027_009690: CJ027_009735 | 10.8 |
| **PPL116** | NRPS-like | PIK34_07725: PIK34_07820 | 22.2 |
| **PPL116** | RiPP-like | PIK34_12060: PIK34_12105 | 10.8 |
| **PPL116** | NRPS | PIK34_21010: PIK34_21020 | 15.6 |
| **PPL116** | NRPS | PIK34_21025 | 9.5 |
| **PPL116** | NRPS | PIK34_21110 | 4.9 |
| **CFBP4641** | NRPS | O2N62_03110: O2N62_03175 | 22.2 |
| **CFBP4641** | RiPP-like | O2N62_04900: O2N62_04950 | 10.9 |
| **CFBP4641** | NRPS | O2N62_20825: O2N62_20835 | 12.8 |
| **CFBP4641** | NRPS | O2N62_20920 | 8.4 |
| **CFBP4641** | NRPS | O2N62_20945 | 5.1 |
| **PPL405** | RiPP-like | O8W29_05580: O8W29_05635 | 10.8 |
| **PPL405** | NRPS | O8W29_12135: O8W29_12230 | 32.3 |
| **PPL405** | NRPS | O8W29_21210 | 4.6 |
| **PPL405** | NRPS-like | O8W29_21220 | 1.4 |
